# Biological and genomic characterization of a new chlorovirus isolate expands the diversity and complexity of giant algal viruses in Brazil

**DOI:** 10.1101/2025.07.24.666624

**Authors:** Isadora Luiza de Jesus Canto, Clécio Alonso da Costa Filho, Lethícia Rodrigues Henriques, Amanda Haisi, João Pessoa Araújo Júnior, Rodrigo Araújo Lima Rodrigues

## Abstract

A novel giant virus was isolated from a freshwater sample collected in Pará de Minas city, Minas Gerais state, Brazil. The new isolate, named BR-AMG2, is a chlorovirus infecting the microalga *Chlorella variabilis*. Members of the family *Phycodnaviridae*, the chloroviruses have large linear dsDNA genomes (285–410 kbp), encoding hundreds of proteins and tRNAs. Despite their ecological and evolutionary importance, chloroviruses remain poorly studied, especially in South America. This study aimed to expand the understanding of this viral group by isolating and fully characterizing new chloroviruses in Brazil. After inoculating alga cultures with environmental samples, cytopathic effects, marked by lysis plaque formation, were observed 72 hours post-sampling. Chlorovirus BR-AMG2 remained stable across a temperature range of - 30 °C to 37 °C, yet was highly sensitive to UV exposure. This virus has a 160 nm icosahedral capsid and establishes a large viral factory in the host cytoplasm. Whole genome sequencing revealed a 338,043 bp genome, containing 11 tRNAs and 388 protein-coding sequences. We observed an extensive DNA methylation machinery, accounting for ∼5% of gene content in this isolate, suggesting a highly methylated genome. Phylogenetic reconstruction and Average Nucleotide Identity analyses confirmed that BR-AMG2 is a new member of the species *Chlorovirus americanus*, the first isolate recovered from a tropical region of the planet. Additionally, we observed a slight expansion of the species pangenome, suggesting ongoing genetic innovation with new isolates. Altogether, our results expand our knowledge about the viral diversity in Brazil and provide valuable insights into the biology of giant algae-infecting viruses.

## INTRODUCTION

Aquatic ecosystems are rich in biodiversity, particularly in the diversity and abundance of viruses, with estimates indicating more than 10^31^ viral particles around the planet (1). Among the most abundant groups in these environments are the so-called giant viruses, especially those associated with protists (2). The term giant virus was first proposed in the late 1990s to describe a virus that infects Chlorella-like algae, known as Paramecium bursaria Chlorella virus 1 (PBCV-1) (3). This virus has a complex pseudoicosahedral capsid approximately 190 nm in diameter, composed of multiple proteins and an internal lipid membrane that encloses a linear double-stranded DNA (dsDNA) genome (4). Until the early 21^st^ century, it was considered the virus with the largest genome and the greatest number of genes known in the virosphere, with 331 kbp and over 400 genes, earning it the designation of a giant virus. This concept was significantly expanded after the discovery of mimiviruses and other giant amoeba-infecting viruses, such as pandoraviruses, and is now commonly used to refer to members of the phylum *Nucleocytoviricota* (5, 6).

The discovery and subsequent extensive studies of PBCV-1 led to the establishment of the family *Phycodnaviridae*, which includes several giant viruses that infect algae. The most studied group within this family is the genus *Chlorovirus* (7). Chloroviruses infect algae in the family Chlorellaceae, which establish symbiotic relationships with protists (e.g., protozoa and heliozoans) and metazoans (e.g., cnidarians), resulting in a complex network of ecological interactions in freshwater environments (8–10). These viruses replicate their genomes in the host cell nucleus and form large viral factories in the cytoplasm, where new particles are assembled and later released through cell lysis (11). The genomes of currently known chloroviruses range from 285 to 410 kbp, encoding hundreds of proteins and tRNAs, which form an islet in the central region of the genome (12, 13). Phylogenomic analyses have identified three groups of chloroviruses, along with several species yet to be officially classified by the International Committee on Taxonomy of Viruses (ICTV), which we refer to as Alphachlorovirus, Betachlorovirus, and Gammachlorovirus (12). This classification aligns with the traditional grouping of chloroviruses based on their host specificity: *Chlorella variabilis* NC64 and *C. variabilis* Syngen 2-3 (Alphachlorovirus); *Micractinium conductrix* Pbi (Betachlorovirus); and *Chlorella heliozoae* SAG3.83 (Gammachlorovirus) (12, 14).

Chloroviruses have been isolated from various locations around the world, particularly in the USA and Europe, but little is known about their presence and diversity in the Global South. Genomic analyses of 36 isolates from different chlorovirus groups revealed an open pan-genome, suggesting there is still much to uncover regarding the genetic and biological diversity of these viruses (15). Brazil contains approximately 12% of the planet’s surface freshwater, yet studies on giant algal viruses in the country remain scarce. In this study, we explored Brazilian aquatic environments in search of chloroviruses, aiming to expand our understanding of their biology and genomics in the Southern Hemisphere and to gain deeper insights into the diversity and evolution of these ecologically important viruses.

## MATERIALS AND METHODS

### Host and virus acquisition

For the viral prospecting study, we used microalgae from the species *Chlorella variabilis* NC64A, *Chlorella variabilis* Syngen 2-3, *Chlorella heliozoae* SAG3.83, and *Micractinium conductrix* Pbi. *Chlorella* species were cultured in modified Bold’s Basal Medium (MBBM) supplemented with tetracycline at 10 µg/mL, while *Micractinium conductrix* was maintained in FES medium. Subcultures were performed weekly at a 1:10 ratio. Cultures were incubated at 25 °C under continuous light in temperature-controlled incubators. The chlorovirus Paramecium bursaria Chlorella virus 1 (PBCV-1) was used as a viral control in the experiments. All algal strains and PBCV-1 were kindly provided by Professor James Van Etten from the Nebraska Center for Virology, University of Nebraska–Lincoln, USA.

### Virus isolation

Environmental water samples were collected from various regions of Brazil, primarily in the state of Minas Gerais, by collaborators from the Virus Laboratory at the Institute of Biological Sciences, Federal University of Minas Gerais, Brazil. All samples were kindly provided for this project and stored at 4°C for later use. Samples were filtered through 0.45 µm nitrocellulose membrane filters, and 1 mL of each was used to infect cultures of the selected microalgae, all in the exponential growth phase at a concentration of 1×10^8^ cells/mL. To confirm the presence of viral infection, the cytophatic effect in alga cultures were observed and then plaque assays were performed on Petri dishes using the double-layer method. Each plate received a bottom layer of 15 mL of medium supplemented with 1.5% agar, followed by a top layer containing 6 mL of medium with 0.75% agar, 1 mL of the sample, and 1 mL of host cells at a concentration of 1×10^9^ cells/mL. Plates were incubated for up to 14 days, monitoring for the appearance of lysis plaques.

From a positive sample, a single lysis plaque was collected and transferred to a microtube containing 1 mL of PBS. The tube was vortexed and centrifuged, and 100 µL of the supernatant was used to inoculate 5 mL of *Chlorella variabilis* NC64A in the exponential growth phase. The infection was incubated on a shaker at 150 rpm for 7 days. After incubation, the culture was centrifuged, and 100 µL of the supernatant was serially diluted and titrated using the double-layer method in 6-well plates. Once the viral titer was determined, a low multiplicity of infection (MOI 0.01) was carried out in 400 mL of *Chlorella variabilis* NC64A (1×10^8^ cells/mL), maintained under constant light and agitation at 25°C for 7 days. The resulting lysate was centrifuged at 4,000 × g for 10 minutes, and the supernatant was filtered through a 0.22 µm membrane filter and ultracentrifuged at 24,000x g for 60 minutes at 4°C. The viral pellet was resuspended in 5 mL of Tris-HCl buffer (pH 7.2) and subsequently purified by centrifugation in a 10– 40% sucrose gradient at 36,000x g for 20 minutes. The purified viral band was collected, and the viral titer was determined using the double-layer assay in 6-well plates.

### Viral stability assays

The isolated virus was subjected to tests to assess its stability under different temperatures and after exposure to ultraviolet (UV) radiation. For the temperature assays, 100 µL aliquots of the purified virus were placed in microtubes and exposed to various temperatures for 10 minutes. The −80 °C and −30 °C treatments were conducted using ultralow freezers; samples at 4°C were stored in a refrigerator, while those at 37 °C, 56 °C, and 100 °C were incubated in a thermoblock. For UV radiation assays, 6-well plates containing 100 µL of purified virus per well were exposed to a UV light source, 15 cm distant, inside a biological safety cabinet. Viral samples were collected at different time points (1, 2, 3, 4, 5, and 10 minutes). In both assays, the treated viral samples were serially diluted and titrated using the double-layer plaque assay in 6-well plates. All experiments were done in biological duplicates.

### Viral replication and host range assays

To assess the growth profile of the viral isolate, particularly in comparison to the model virus PBCV-1, infections at an MOI of 0.01 were performed using 3.6×10^8^ *Chlorella variabilis* NC64A cells. Infections were maintained on a shaker at 150 rpm for 24 hours. After incubation, cultures were centrifuged at 4,000x g for 10 minutes, and the supernatant was collected, serially diluted, and titrated using the double-layer plaque assay in 6-well plates.

We also evaluated host range of the isolated virus. For this, 10 µL of purified virus was added to Falcon tubes containing 5 mL of microalgal cultures from the species *Chlorella heliozoae, Micractinium conductrix*, and *Chlorella vulgaris*. Tubes were maintained under agitation for 7 days, and the cultures were monitored daily for potential cytopathic effects.

### Transmission Electron Microscopy analysis

The purified viral isolate was analyzed at the UFMG Microscopy Center for transmission electron microscopy (TEM) to assess the morphology and replication cycle. For negative staining, 100 µL of the viral suspension was treated with osmium tetroxide and placed onto copper grids for observation. Additionally, embedded samples were prepared from an infection at MOI 1 in 5 mL of *Chlorella variabilis* NC64A culture (total of 5×10^8^ cells). The infection was maintained on a shaker at 150 rpm for 8 hours. After this period, the culture was centrifuged at 4,000×g for 10 minutes, and the pellet was resuspended in 1.5 mL of Karnovsky solution, sonicated briefly, and kept under agitation at 150 rpm for 4 hours. The material was then washed three times with 0.1 M phosphate buffer (pH 7.3) and sent to the Microscopy Center, where it was post-fixed with osmium tetroxide, embedded in resin, sectioned, and contrasted. Both preparations were visualized using a Tecnai G2-12-Spirit BioTwin FEI 120 kV transmission electron microscope. Images were processed and analyzed using the ImageJ software.

### Genome sequencing, assembly, and annotation

The purified virus was sequenced using the Illumina MiSeq i100 with a paired-end 2×300 bp library, prepared with the Illumina DNA Prep kit (Illumina Inc., San Diego, CA, USA). Raw reads were analyzed using FastQC for quality assessment and trimmed to remove low-quality bases with Trimmomatic (16). Assembly was performed using the SPAdes program (version 3.15.5) with default parameters (17), implemented in Galaxy (https://usegalaxy.eu/). The resulting scaffold was compared to sequences from the National Center for Biotechnology Information (NCBI) database using BLASTn and organized with MeDuSa online (18). Open reading frames (ORFs) were predicted using Prodigal (version 2.6.3) (19), and tRNAs were identified using the default parameters of ARAGORN (20). ORFs were annotated using BLASTp (NCBI nr database), HHpred, and InterProScan, following the strategy described elsewhere (12). Functional categorization of the predicted proteins was performed based on Orthologous Groups of Nucleo-Cytoplasmic Viruses (21).

### Genomic and phylogenetic analysis

The genome was visualized in circular format using Proksee (22). Average Nucleotide Identity (ANI) analysis was performed with FastANI (23), using the complete genomes of eight chlorovirus isolates. Synteny analysis was conducted using DiGAlign (24). Clusters of orthologous groups of genes (COGs) were constructed with OrthoFinder using default parameters (25). Methyltransferases COGs were constructed using ProteinOrtho6 with the following parameters: e=1e-5; cov=50; identity=50 (26). Pan-genome evolution was assessed based on the stepwise addition of isolates, and the COG-sharing pattern was visualized using Gephi 0.10 (27). Pan-genome and methyltransferase-related networks were generated using ForceAtlas, a force-directed layout algorithm implemented in Gephi, and were minimally edited manually to improve the visualization of singleton nodes.

For phylogenetic reconstruction, amino acid sequences of the chlorovirus DNA polymerase B family genes were aligned using MUSCLE with default parameters (28) and subsequently trimmed with TrimAl to remove misaligned regions (29). The phylogenetic tree was constructed using IQ-TREE2 with the maximum likelihood method and 1,000 bootstrap replicates (30), employing the best-fit evolutionary model selected by ModelFinder (31). The resulting tree was visualized using iTOL v6 (32).

## RESULTS

### A new chlorovirus found in the waters of Minas Gerais, Brazil

Water samples collected in September 2024 were filtered and inoculated into microalgae cultures for up to 14 days to observe cytopathic effects. In a sample collected from a lake in Bariri Municipal Park, in the city of Pará de Minas, Minas Gerais state (Figure 1A), we observed the presence of lysis plaques in *C. variabilis* NC64A after 72 hours of infection, indicating viral activity. The lysis plaque was collected and inoculated into a new algal culture for viral propagation and subsequent rounds of plaque purification. We named the new isolate BR-AMG2. Once purified, a fresh algal culture was infected, and complete cell lysis and total culture clearance were observed within 7 days post-infection, an effect similar to that of PBCV-1, the prototype *Chlorovirus* isolate (Figure 1B). The BR-AMG2 isolate forms lysis plaques of approximately 2 mm in diameter after 7 days of infection (Figure 1B), which is considered medium-sized when compared to other known *Chloroviruses* (33). Transmission electron microscopy revealed an icosahedral particle approximately 160 nm in diameter, with an internal membrane surrounding the genome (Figure 1C). Viral particles attach to the cell wall and upon entering the host cell, BR-AMG2 establishes a large viral factory in the cytoplasm, where new viral particles are assembled and later released through cell lysis (Figure 1D). After 24 hours of infection, we observed a ∼175-fold increase in BR-AMG2 particles in the supernatant, slightly lower than the yield observed for PBCV-1 (Figure 1E). The new isolate demonstrated good thermal stability between -30 °C and 37 °C, maintaining titers above 10^9^ PFU/mL. However, the titer dropped by 2 logs when stored at -80°C and by approximately 5 logs after exposure to 56 °C for 10 minutes (Figure 1F). The virus was completely inactivated at 100 °C. When exposed to UV radiation, we observed a gradual decrease in viral titer between 0 and 5 minutes of exposure, dropping from 3.4×10^9^ PFU/mL to 1×10^3^ PFU/mL, a 99.9999% reduction in infectious particles (Figure 1G). Complete inactivation occurred after 10 minutes of UV exposure.

**Figure 1:**
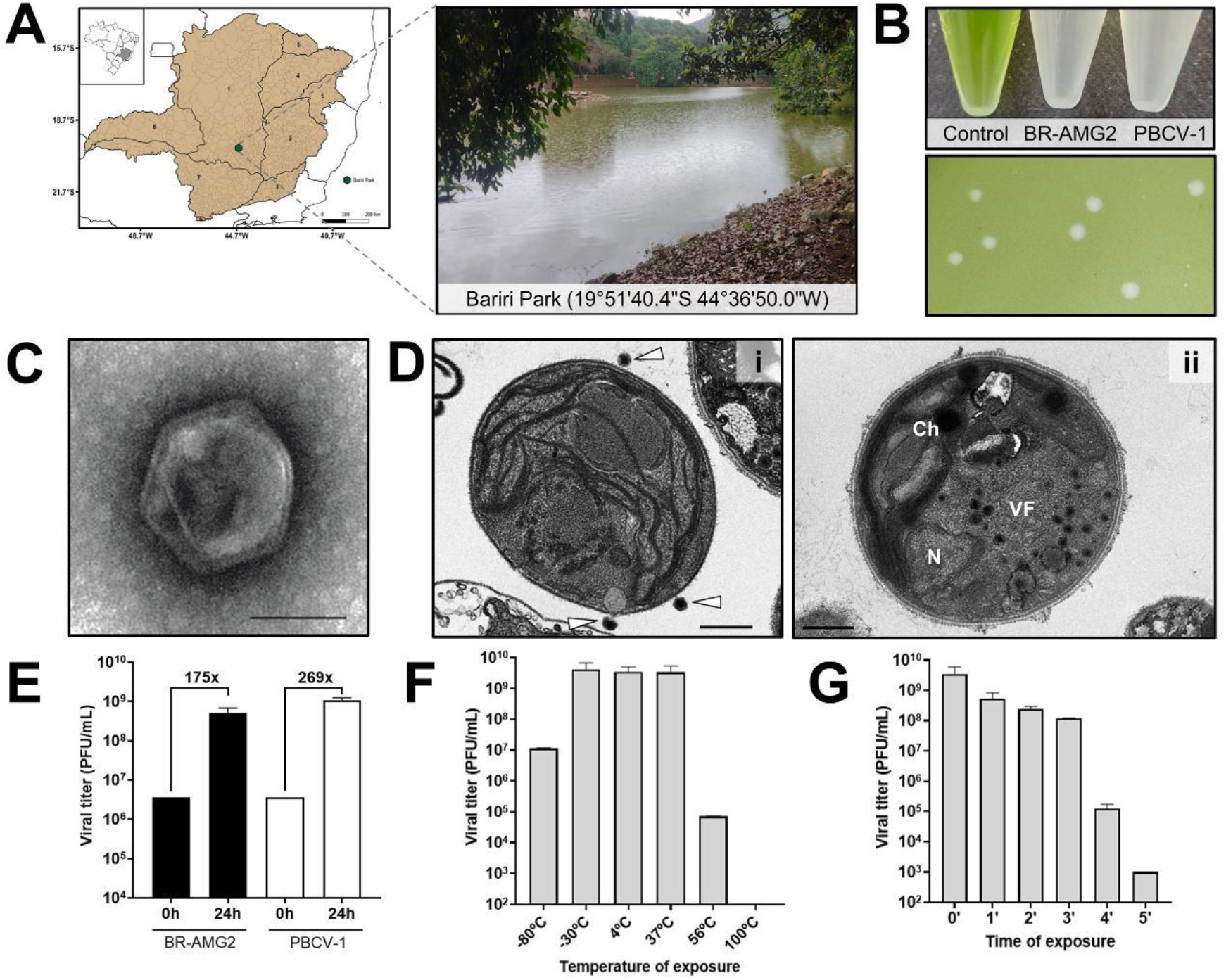
Isolation and biological characterization of Chlorovirus BR-AMG2. A) Isolation site of a new chlorovirus. The map was created with QGIS software and is divided based on the main river basins present in the state, named (1) São Francisco River; (2) Pardo River; (3) Jequitinhonha River;(4) Mucuri River; (5) Doce River; (6) Paraíba do Sul River; (7) Paraíba River; (8) Grande River; and (9) Piracicaba River. The area highlighted in green indicates the Bariri Municipal Park, located within the São Francisco River basin, where the BR-AMG2 was isolated; B) Infected cultures of chloroviruses and plaque morphology of BR-AMG2. Plaques are average 2 mm in diameter; C) Negative staining electron microscopy of BR-AMG2, revealing an icosahedral particle with inner membrane; D) Transmission electron microscopy images of *Chlorella variabilis* NC64A infected with BR-AMG2 exhibiting (i) a cell being infected – white arrows point to viral particles, and (ii) a large viral factory in the cell cytoplasm. N: Nucleus, Ch: Chloroplast, VF: Viral factory; E) Viral yield within 24 hours of infection; F) Temperature resistance of BR-AMG2; G) UV radiation sensitivity of BR-AMG2. Biological assays were performed in duplicate. Scale bars: 100 nm (C); 500 nm (D).

### The Brazilian isolate is a new member of the species *Chlorovirus americanus*

The genome of the new isolate was sequenced using the Illumina paired-end method, generating a total of 914,000 reads with quality scores above Q30. *De novo* assembly resulted in seven scaffolds with an average coverage of 125X. BLASTn analysis identified *Chlorovirus americanus* isolates (e.g., AR158, MA-1D, and NY2A) as the best hits. We used these references to organize the scaffolds, resulting in a final genome of 338,043 bp with a GC content of 40.4%. Based on the DNA polymerase B family gene, a well-established phylogenetic marker for viruses in the phylum *Nucleocytoviricota* (6), we confirmed that BR-AMG2 clusters with other *Chlorovirus americanus* isolates (subgenus Alphachlorovirus) with strong statistical support (bootstrap = 100) (Figure 2A).

**Figure 2:**
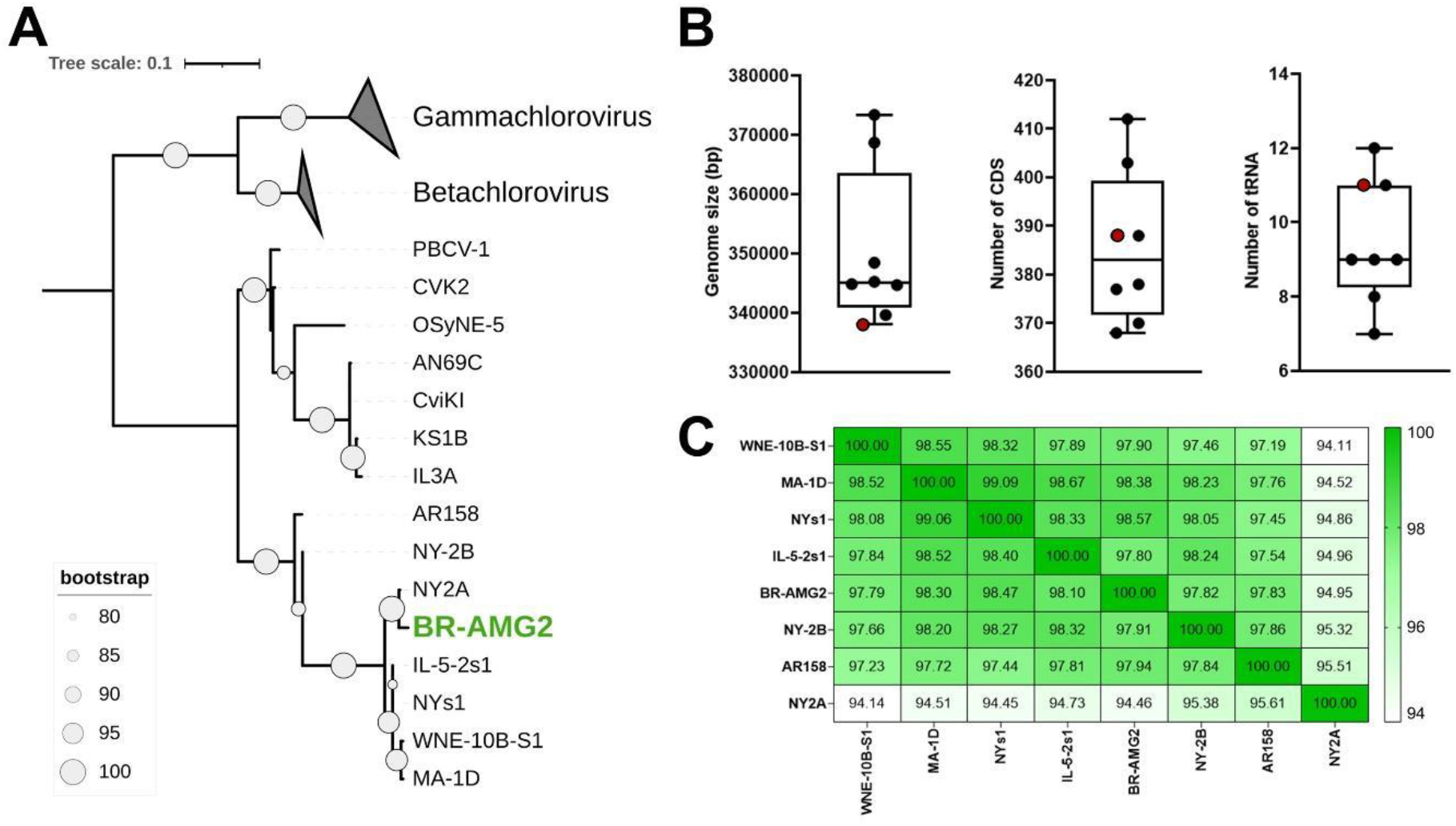
Chlorovirus BR-AMG2 phylogeny and genomic features. A) Phylogenetic tree based on DNA polymerase B family gene, including all three groups of chloroviruses, evidencing the position of isolate BR-AMG2 (green label). The tree was rooted using Betachloroviruses and Gammachloroviruses as outgroups. Scale bar represents rate of amino acid substitution; B) Boxplots of general genomic features of *Chlorovirus americanus* isolates. The BR-AMG2 is highlighted as red dots; C) Heatmap showing ANI values including all eight *Chlorovirus americanus* isolates. Scale bar indicates ANI values ranging from 94 to 100%.

BR-AMG2 has the smallest genome reported within its species, whose genome sizes average approximately 343 kbp, ranging from 338 kbp to 373 kbp (e.g., WNE-10B-S1) (Figure 2B; Table S1). A total of 388 coding sequences (CDSs) longer than 40 amino acids were predicted, matching the number in isolate AR158 and slightly above the species average (373 CDSs). Interestingly, isolate NY2A has the highest number of predicted genes (n = 412), despite having a genome smaller than the species average (368 kbp). Although BR-AMG2 has the smallest genome among known *C. americanus* isolates, it encodes more genes than half of them (Figure 2B; Table S1).

Additionally, 11 tRNAs were predicted for BR-AMG2, above the species average of 8 tRNAs and equal to WNE-10B-S1 and MA-1D (Figure 2B; Table S1). Average Nucleotide Identity (ANI) analysis revealed values above 94% shared with other *C. americanus* isolates, corroborating the phylogenetic placement of BR-AMG2 as a new member of the species (Figure 2C). Among the isolates, NY2A was the most divergent, sharing an average of 95% identity with the others, whereas BR-AMG2 showed approximately 98% identity with its conspecifics. When compared to PBCV-1 (*Chlorovirus vanettense*), BR-AMG2 shared an average genomic identity of 85.8%.

### Genomic analyses of BR-AMG2 isolate reveal an extensive DNA methylation machinery in *Chlorovirus americanus*

The genome of the BR-AMG2 isolate has a coding density of 90.8%, with 388 coding sequences (CDSs) homogeneously distributed between the forward and reverse strands. Genes have an average GC content of 40.8%, ranging from 24.8% to 58.4%, and we found 11 tRNAs, all located on the reverse strand (Figure 3A). In general, chloroviruses exhibit a tRNA islet positioned near the center of the genome (13). Interestingly, in BR-AMG2, the tRNA cluster is slightly shifted to the right, located between positions 191,717 and 192,847. We observed a similar pattern in isolate MA-1D (genome size = 339,653 bp). In the other *Chlorovirus americanus* isolates, the central genome positioning of the tRNA islet is maintained (Table S2). Among the 11 tRNAs encoded by BR-AMG2, eight amino acids are represented, including one duplication each for asparagine, leucine, and lysine, the latter two featuring distinct anticodons (Table S2). The tRNA repertoire of BR-AMG2 is similar to that of other members of the species, with no unique tRNA identified.

**Figure 3:**
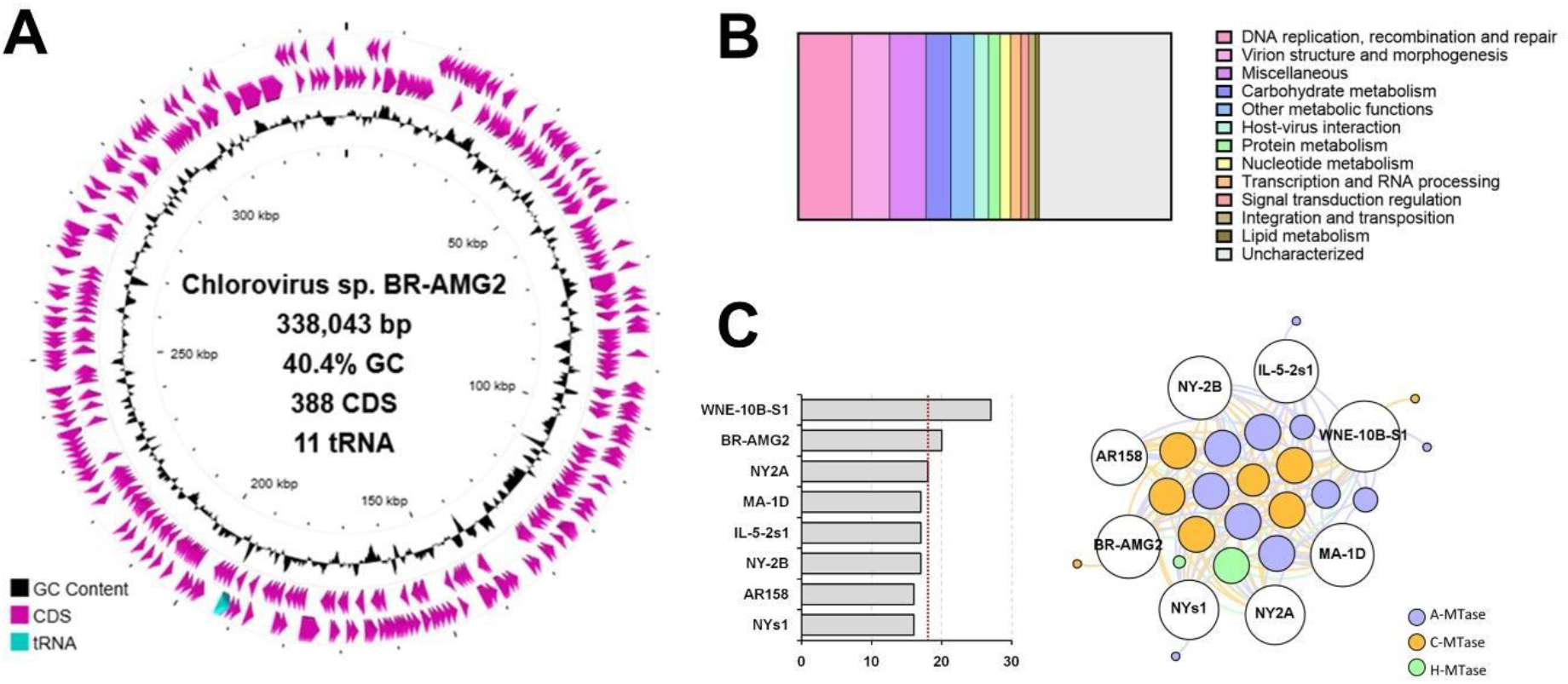
Genome annotation of Chlorovirus BR-AMG2. A) Genomic map of the isolate BR-AMG2, showing the CDSs in pink, tRNA in cyan, and GC content distribution in black ring; B) Functional categorization of genes predicted in BR-AMG2 genome; C) Abundance of methyltransferases and gene-sharing within isolates of Chlorovirus americanus. Red line in the column chart represents the mean value of genes found in the viral genomes. In the network, the node sizes are proportional to the degree of connection. White nodes represent the viruses and are labeled with the isolates’ names, while colored nodes represent different methyltransferases.

Of the 388 predicted CDSs, over one-third (n = 137; 35.3%) have no known function (Figure 3B). Three of these are ORFans, with no detectable similarity to any known protein sequences in public databases. The identification of such novel genes, albeit limited in number, suggests the existence of unexplored genetic diversity among chloroviruses. Among the annotated genes, we identified a wide array of enzymes involved in carbohydrate metabolism, including glycosyltransferases, chitinases, chitosanases, and alginate lyases, which have also been reported in other chloroviruses. Additionally, genes related to lipid and protein metabolism were found, including lipid hydrolases and E3 ubiquitin ligases, enzymes responsible for the degradation of lipids and proteins, respectively. A total of 11 genes involved in transcriptional regulation or RNA modification were identified, including transcription factors (e.g., TFIIB, TFIID, and TFIIS), RNase III, and an mRNA capping enzyme. No RNA polymerase subunit was detected, consistent with observations in other chloroviruses (12, 14, 34). Numerous genes involved in virion structure and morphogenesis were also identified (Figure 3B), including the major capsid protein (Vp54), several minor capsid proteins homologous to those found in PBCV-1 (4), and the DNA packaging ATPase, a gene conserved across the phylum *Nucleocytoviricota* and implicated in genome encapsidation (35, 36).

Among the functional categories, the most abundant was DNA replication, recombination, and repair, with 56 genes (14.4%) involved in these processes (Figure 3B). It includes genes commonly found in giant viruses, such as DNA polymerase, DNA topoisomerase, DNA primase, and helicases, as well as various endonucleases, including those with GIY-YIG catalytic domains, enzymes previously described in other chloroviruses. Notably, we observed a large number of methyltransferases (MTases) dispersed throughout the BR-AMG2 genome. A total of 20 enzymes involved in methylation of adenine, cytosine, and histones were identified, accounting for approximately 5% of the isolate’s gene content. MTases were also found in other isolates of *C. americanus*, with an average of 18 MTase genes per genome (Figure 3C). Only isolate WNE-10B-S1 harbors more MTases than BR-AMG2, with 27 genes distributed across its genome. Proportionally, we did not observe major differences among the isolates, with MTase-related genes comprising between 4.12% and 6.70% of their total gene content.

Most of the MTases are orthologous, forming conserved gene clusters among the isolates. However, we identified unique MTases in four isolates, including a cytosine-specific DNA methyltransferase in BR-AMG2 (Figure 3C). Although MTases are also present in chloroviruses from other subgroups, the average number observed in *C. americanus* is higher than in other species, including various alphachloroviruses (data not shown).

### Genomic conservation and pangenome evolution of *Chlorovirus americanus*

Synteny analysis revealed a high conservation of gene organization among *Chlorovirus americanus* isolates, with a strong degree of gene identity, reinforcing the results observed in the ANI analysis (Figure 4). A genomic region reorganization in isolate BR-AMG2 compared to the others can be observed near the 190 kbp region. Minor variations are also present in the genomes of isolates IL-5-2s1 and MA-1D, both maintaining a high degree of identity (>95%). Although isolate NY2A shows a lower overall similarity to the other isolates, its genomic organization remains well conserved (Figure 4). The terminal regions of the genomes are less conserved relative to the rest of the genome, exhibiting approximately 85% identity in some isolates (e.g., BR-AMG2 and NY-2B).

**Figure 4:**
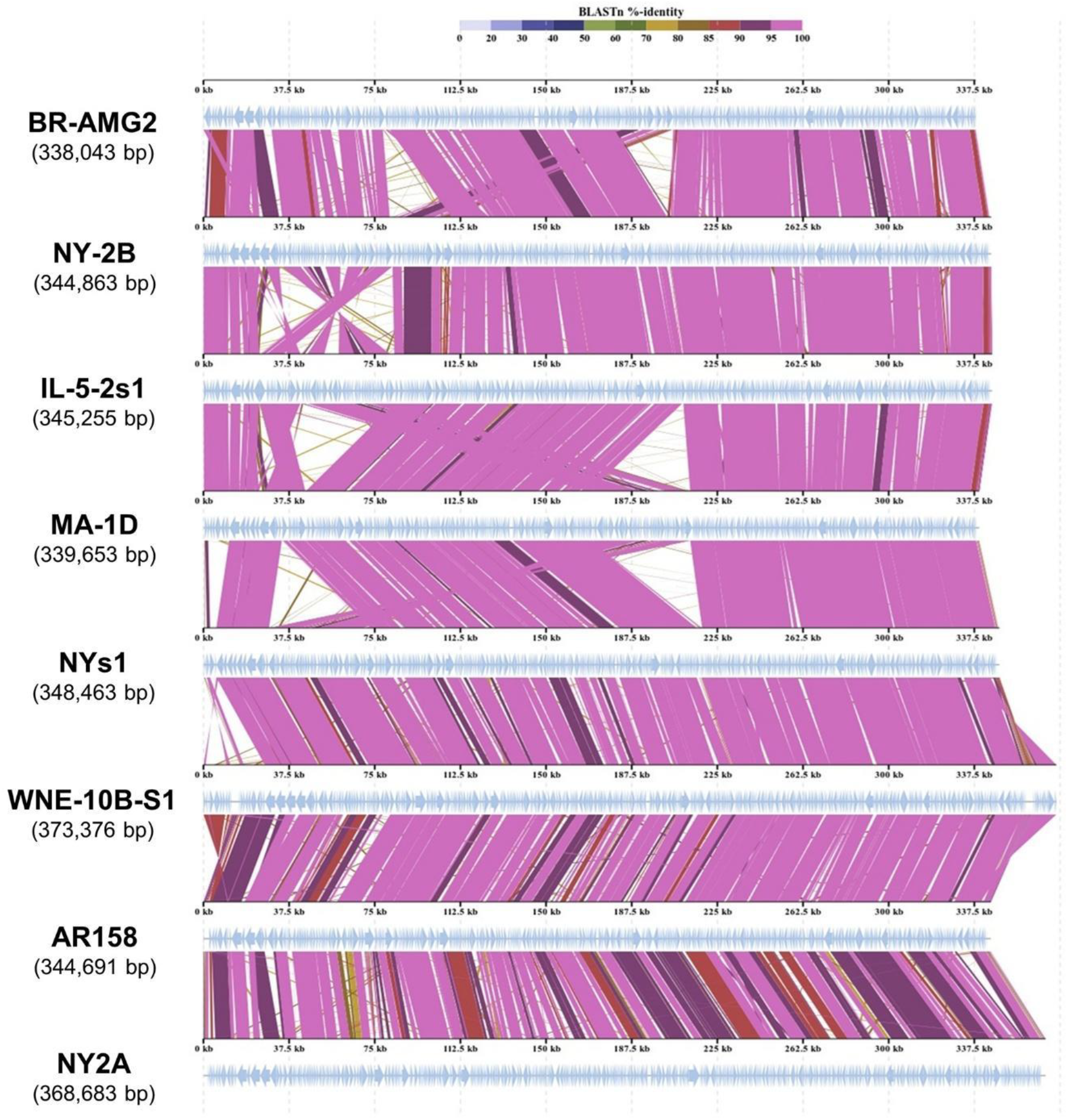
Genome synteny of Chlorovirus americanus. Viral genomes were aligned, and colored lines represent BLASTn identity values. The name of the isolates and their genome sizes are indicated on the left side of the respective genome representation.

After defining the species to which the Brazilian isolate belongs, we evaluated its impact on the evolution of the *Chlorovirus americanus* pangenome and the pattern of clusters of orthologous genes (COGs) shared among viral isolates (Figure 5). A total of 3,162 predicted genes from the eight isolates were grouped into 445 COGs, constituting the species’ pangenome. As we added isolates, we observed a marginal increase in COGs, ranging from 1.15% to 6.61% depending on the isolate, reaching a cumulative increase of 24.64% (Figure 5A). Conversely, we observed a slightly more pronounced reduction in shared COGs among the isolates (core genome), followed by stabilization of the curve (Figure 5A). Comparing the first two isolates analyzed, NY2A and NY-2B, there was an 8.4% reduction in the core genome. The addition of the remaining isolates resulted in a mild decrease, between 2.1% and 0%, indicating stabilization. Overall, we calculated a 12.9% reduction in the core genome of *Chlorovirus americanus* considering all isolates of the species. The inclusion of BR-AMG2 contributed to a 1.59% increase in the number of COGs and only a 0.32% decrease in the core genome. Of the 445 COGs identified, 311 (69.9%) are shared by all isolates, constituting the core genome; 99 (22.2%) are shared by two or more isolates, constituting the accessory genome; and 35 (7.9%) are singletons (Figure 5B). Our analysis identified four singletons unique to the new Brazilian isolate, a number comparable to other isolates, except for NY-2B, which had no unique COGs, and NY2A, which contained 10 unique COGs (Figure 5B).

**Figure 5:**
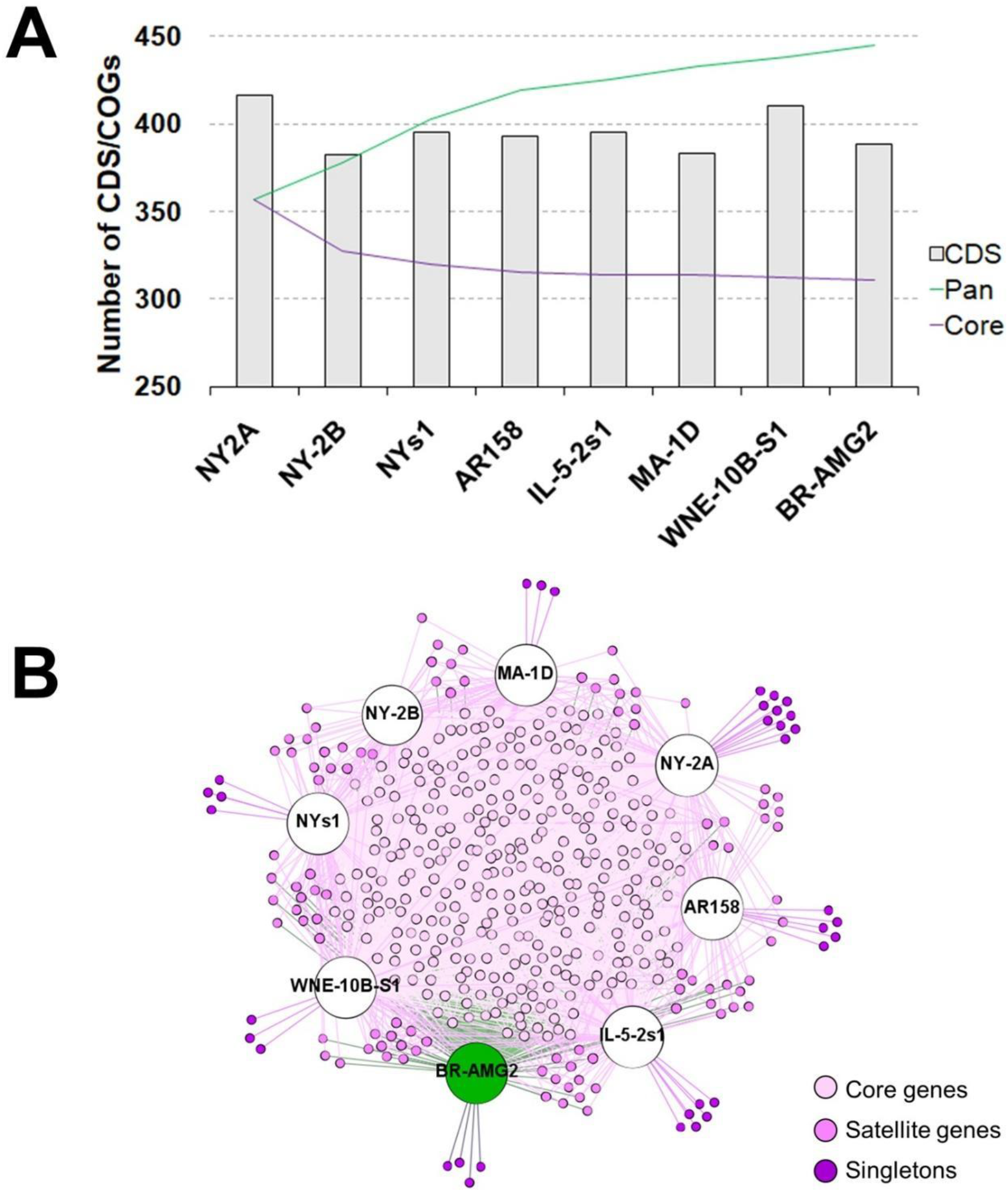
Pan-genome evolution and COG-sharing pattern of Chlorovirus americanus. A) Pan-genome evolution including all eight isolates that belong to the species Chlorovirus americanus. Columns indicate the amount of CDS predicted in each genome. Green curve indicates the increase in the pan-genome, and purple curve refers to the decrease in the core genome with the addition of new isolates to the species; B) Bipartite network including viruses (BR-AMG2 is indicated with green node, while other isolates are represented as white nodes) and the shared COGs (colored nodes). The node sizes are proportional to the degree of connection.

## DISCUSSION

Chloroviruses were discovered over 40 years ago, and much of our current knowledge is based on studies of PBCV-1, which for years was considered the prototype of the genus *Chlorovirus* [for a detailed review, see Van Etten et al., 2020 - (37)]. The isolation of new viruses has enabled progressive advances in understanding the diversity and evolution of this group of aquatic viruses, revealing several species that cluster into three well-defined phylogenetic clades (12, 34). These viruses have been isolated from freshwater samples in various regions worldwide; however, the full extent of chlorovirus genomic and biological diversity remains far from fully characterized (12, 14, 34, 38). In Brazil, only two isolates have been described to date: one alphachlorovirus and one gammachlorovirus (14, 39). Given Brazil’s renowned biodiversity and its possession of over 10% of the planet’s surface freshwater, there is a significant knowledge gap regarding the presence, diversity, and ecology of chloroviruses both in Brazil and globally. In this work, we contribute to filling this gap by isolating and characterizing a new chlorovirus from the waters of Minas Gerais, one of Brazil’s largest states.

The BR-AMG2 isolate presents a hexagonal particle approximately 160 nm in diameter, slightly smaller than that described for other chloroviruses (4, 40). We expect that its structure is highly similar to that of PBCV-1, as we detected homologs of several structural proteins, including the major capsid protein (MCP Vp54 and its variants MCPv1 and MCPv5), the penton protein (P1) and its variant (P1v1), as well as 19 of 22 minor capsid proteins (4). The three missing proteins (P20, P21, and P23) may be present but remain undetected due to high divergence beyond our search criteria. Like other chloroviruses, BR-AMG2 enters the host cell by inducing a pore in the cell wall, possibly facilitated by alginate lyase and other glycosyl hydrolases present in its particle (41, 42). The virus establishes a large viral factory that occupies nearly the entire cytoplasm, where new particles are assembled and subsequently released through cell lysis (11). We estimate that each infected cell releases between 100 and 200 particles, a burst size comparable to other chloroviruses (37). We observed no significant alterations in chloroplasts of infected cells, suggesting that this organelle is not exploited during viral replication. Additional studies should be done to evaluate the impact of cell organelles in chloroviruses replication. Interestingly, BR-AMG2 is resistant to temperature variations between -80 °C and 56 °C during 10-minute exposures. However, prolonged freezing caused a significant decline in viral infectivity, indicating that, like PBCV-1, it cannot tolerate extended exposure to very low temperatures (37). Additionally, the virus is highly sensitive to UV radiation, being completely inactivated after only 10 minutes of exposure. Unlike some other giant viruses (43), BR-AMG2 lacks a robust DNA repair machinery, which may explain its low resistance to UV radiation. Additional studies exposing chlorovirus-infected cells to UV radiation may provide insights into this feature.

BR-AMG2 is the first Brazilian isolate, and the second one found outside the USA, classified as *Chlorovirus americanus*, an alphachlorovirus. Both phylogenetic and ANI analyses support this classification, following established strategies used for dozens of other chloroviruses (12, 34). Although BR-AMG2 has the smallest genome of the species, its structural features do not significantly differ from other known chloroviruses. However, analysis of its genetic content revealed a substantial number of genes encoding methyltransferases, which constitute more than 5% of its gene repertoire, suggesting a highly methylated genome. Methyltransferases have been detected in other chloroviruses, and liquid chromatography analyses have shown wide variation in methylation levels across viral genomes, with the NY2A isolate exhibiting up to 45% methylated cytosines and 37% methylated adenines (44). We identified 18 MTases in NY2A and 20 in BR-AMG2, suggesting a comparable methylation level in the Brazilian isolate. The extent of methylation in BR-AMG2 and other *Chlorovirus americanus* isolates remains to be determined to establish whether this is a species-wide or an isolate-specific trait. For PBCV-1 (*Chlorovirus vanettense*), SMRT sequencing data evidenced that viral adenines are stably methylated (45), and earlier studies suggested at least 2% of cytosines and adenines are methylated (46). Similar methylation levels have also been reported in some betachloroviruses, with 5-methylcytosine (5mC) ranging from 14% to 43% and N6-methyladenine (6mA) up to 18% (47). Biochemical characterization of some chlorovirus MTases confirmed that these genes are expressed and that the enzymes methylate the viral genome (48, 49). DNA methylation regulates various biological processes, including gene expression and, in viruses, protection against host defenses (50, 51). In chloroviruses, some MTases are accompanied by restriction endonucleases, forming a restriction-modification system (52). Further study of BR-AMG2’s MTases, especially its unique cytosine-specific methyltransferase, may provide valuable insights into virus-host interactions.

The isolation and characterization of new viruses have uncovered unique features of chloroviruses and fostered progress across virology, biochemistry, and viral biotechnology. Notably, several enzymes encoded by chloroviruses have been characterized and developed into molecular biology tools, including endonucleases, methyltransferases and ligases (37, 53). By isolating a new virus from Brazilian waters, we advance knowledge of the presence and diversity of these viruses, particularly in the Southern Hemisphere. The slight expansion of the species’ pangenome suggests ongoing potential for genetic innovation, especially considering the diverse chlorovirus clades (15). Given Brazil’s continental scale, exploration for new viruses in different regions promises significant advances in the study of giant algal viruses.

## ACKNOWLEDGMENTS

We would like to thank all members of the Grupo de Estudo em Phycodnavirus (Gephy) and the Virus Laboratory at UFMG for their collaboration and scientific discussions during the development of this work. We also thank the members of Prof. James Van Etten’s lab at the University of Nebraska-Lincoln for providing cells and viruses, and all the support in establishing protocols and experiments that culminated in the isolation of the novel virus presented here. This project was registered at the SISGEN n° A048F34.

## DATA AVAILABILITY

The genome sequence of Chlorovirus americanus isolate BR-AMG2 was deposited in the NCBI GenBank database with accession number PV975788. The datasets generated and/or analyzed during the current study are available in the GenBank repository.

## SUPPLEMENTARY MATERIAL

**Table S1:** General information about *Chlorovirus americanus* isolates.

**Table S2:** Diversity of tRNA in *Chlorovirus americanus*.

