## Supplementary material for "Biological and genomic characterization of a new chlorovirus isolate expands the diversity and complexity of giant algal viruses in Brazil": Suplementary Table 1

**Table S1:** General information about *Chlorovirus americanus* isolates.

| Virus | Genome size (bp) | CDS count | tRNA count | GC content | Location of isolation | Year of description | GenBank accession |
|---|---|---|---|---|---|---|---|
| **BR-AMG2** | 338043 | 388 | 11 | 40.4% | Minas Gerais, Brazil | 2025 | PV975788 |
| **MA-1D** | 339653 | 368 | 11 | 40.69% | Massachusetts, USA | 2013 | JX997172 |
| **AR158** | 344691 | 388 | 7 | 40.76% | Buenos Aires, Argentina | 2007 | NC_009899 |
| **NY-2B** | 344863 | 370 | 8 | 40.53% | New York, USA | 2013 | JX997182 |
| **IL-5–2s1** | 345255 | 377 | 8 | 40.19% | Illinois, USA | 2013 | JX997170 |
| **NYs1** | 348463 | 378 | 8 | 40.71% | New York, USA | 2013 | NC_043235 |
| **NY2A** | 368683 | 412 | 7 | 40.69% | New York, USA | 2007 | NC_009898 |
| **WNE-10B-S1** | 373376 | 403 | 11 | 40.87% | Nebraska, USA | 2024 | PP681875 |
