## Supplementary material for "Biological and genomic characterization of a new chlorovirus isolate expands the diversity and complexity of giant algal viruses in Brazil": Suplementary Table 2

**Table S2:** Diversity of tRNA in *Chlorovirus americanus*.

| Virus | Islet start (nt)* | Islet end (nt)* | Genome position | Island size (nt) | Leu (taa) | Leu (caa) | Ile (tat) | Asn (gtt) | Arg (tct) | Gly (tcc) | Lys (ctt) | Lys (ttt) | Gln (ctg) | Val (aac) |
|---|---|---|---|---|---|---|---|---|---|---|---|---|---|---|
| **NY-2B** | **169407** | **170240** | **49%** | **833** | 1 | 1 | 1 | 1 | 1 | 1 | 1 | | | 1 |
| **IL-5-2s1** | **175844** | **176677** | **51%** | **833** | 1 | 1 | 1 | 1 | 1 | 1 | 1 | | | 1 |
| **NY2A** | **194698** | **195555** | **53%** | **857** | 1 | 1 | | 1 | 1 | 1 | 1 | | | 1 |
| **NYs1** | **180325** | **182145** | **52%** | **1820** | 1 | 1 | 1 | 1 | 1 | 1 | 1 | | | 1 |
| **AR158** | **172099** | **172781** | **50%** | **682** | 1 | 1 | 1 | 1 | 1 | 1 | | | | 1 |
| **MA-1D** | **134220** | **135350** | **40%** | **1130** | 1 | 1 | 1 | 2 | 1 | 1 | 1 | 1 | 1 | 1 |
| **WNE-10B-S1** | **193976** | **195106** | **52%** | **1130** | 1 | 1 | 1 | 2 | 1 | 1 | 1 | 1 | 1 | 1 |
| **BR-AMG2** | **145197** | **146327** | **43%** | **1130** | 1 | 1 | 1 | 2 | 1 | 1 | 1 | 1 | 1 | 1 |

*Genome coordinates may vary from information available in GenBank due to reorganization in orientation to allow proper comparison.
